# A mechanism of T cell dependent selection of antigen engaged Germinal Center B cells

**DOI:** 10.1101/061879

**Authors:** Vinod Krishna, Kurtis E. Bachman

**Author notes:** Electronic mail; Corresponding author.

## Abstract

A model of B cell affinity selection is proposed, and an explanation of peripheral tolerance mechanisms through antibody repertoire editing is presented. We show that affinity discrimination between B cells is driven by a competition between obtaining T cell help and removal of B cells from the light zone, either through apoptosis or by a return to the dark zone of germinal centers. We demonstrate that this mechanism also allows for the negative selection of self reactive B cells and maintenance of B cell tolerance during the germinal center reaction. Finally, we demonstrate that clonal expansion upon return to the germinal center dark zone amplifies differences in the antigen affinity of B cells that survive the light zone.

## I. INTRODUCTION

The ability of B cells to form antibodies against unknown foreign antigens is fundamental to immunity against infection. B cells are able to synthesize antibodies through an evolutionary process which involves the mutation and selection of their B cell receptors (BCRs) for enhanced antigen-specific recognition, resulting in affinity maturation of B cells. In the initial stage of early antigen engagement, B cells are enriched for those with receptors that have an adequate antigen binding affinity. The enriched B cell populations then migrate to specialized anatomical structures that form in the lymph nodes and similar organs, known as germinal centers (GC), where B cell receptor affinity maturation occurs. B cells in the GC undergo clonal expansion and somatic hypermutation (SHM) at the BCR. This is followed by antigen uptake by the hypermutated B cells from GC resident follicular dendritic cells (FDC’s) and selection between the resulting antigen presenting hypermutated B cells for affinity maturation by follicular helper T cells (Tfh cells).^1^

According to the classic model of GC B cell affinity maturation, GC B cell somatic hypermutation and clonal expansion occur in a spatially distinct GC “dark zone” (DZ), while antigen loading by follicular dendritic cells (FDC’s) and B cell selection occur in the so-called GC “light zone” (LZ) (Fig 1a).^1^ While this model of B cell affinity maturation explains the broad contours of how immunological tolerance is maintained or re-established by the GC reaction, it is not clear how B cell interactions with antigen bound FDC’s and Tfh cells in the GC result in both a positive selection for highly antigen specific BCRs, and a negative selection against self reactive B cells.

Experiments have shown that the affinity selection of B cells in the GC light zone is limited by access to costimulation by Tfh cells.^2–5^ On the other hand, while somatic hypermutation and clonal expansion of B cells result in a few clones with improved antigen affinity, the majority of hypermutated B cells are likely to be either self reactive or have degraded affinity for antigen.^6–8^ In addition, Tfh cells recognize short peptide antigen epitopes through T cell receptor (TCR) binding to pMHC complexes, while affinity maturation requires optimizing the binding affinity of the BCR to the whole antigen. A central question is to reconcile these observations and describe the mechanism that governs the selection of high affinity, antigen specific B cells out of the large pool of hypermutated B cells with low and intermediate affinity, while at the same time also eliminating hypermutated B cells with cross reactivity to both antigen and self proteins. Specifically, in this paper we address how B cells that enter the GC LZ could undergo both a positive selection for antigen binding affinity and a negative selection against autoreactive B cells through encounters with Tfh cells. In addition, we examine how selection of Tfh cell specific antigen epitopes could also result in selection for higher BCR antigen affinity.

In this work, we propose a theoretical model to address these questions, based on the recent observations that a substantial fraction of B cells return to the GC dark zone after encountering cognate Tfh cells,^5,9^ and the property that GC B cells undergo apoptosis in large numbers, with experimental studies implicating apoptosis as an important mechanism for editing out self reactive B cells in the GC.^4,10–12^ We show that antigen binding specificity and negative selection against self antigen can be achieved by a tradeoff between Tfh cell binding and the removal of B cells in the GC light zone, either due to apoptotic clearance or by cycling of B cells back to the GC dark zone due to successful Tfh cell costimulation. We then discuss how apoptosis and B cell cycling out of the LZ during the GC B-Tfh costimulatory reactions greatly increases selection between distinct antigen epitopes presented by the B cell on its surface. Based on the observed link between the amount of antigen bound by a B cell to the amount of T cell specific epitopes presented, we describe how T cell discrimination between different epitopes results in selection for B cells with higher affinity in this framework. Finally, we show how the same mechanisms that govern positive selection for higher antigen affinity can also result in a negative selection against self reactive B cells.

## II. THEORETICAL MODELS OF B CELL AFFINITY SELECTION

At a first glance, B cells are distinguished from each other by Tfh cells by the rate at which a given Tfh cell binds different B cells. This difference in rates is related to the differential amount of antigen processing and peptide presentation by B cells. We assume that individual Tfh-B cell encounters are independent and irreversible, since B-Tfh cell interactions drive internal B cell signaling pathways and alter B cell state, at the very least Tfh cell binding drives anti-apoptotic signaling in B cells.^10,11^ In what follows, we first relate antigen binding affinity and consequent antigen presentation, to make a simple argument to examine the maximal affinity discrimination possible when only BCR antigen affinity and equilibrium interactions with Tfh cells are considered.

If a given antigen produces a maximum of *p* peptides upon internalization after B cell binding and uptake, then when a B cell binds and internalizes *N* copies of the antigen, a maximum of *N*_*p*_ peptides can be expressed on the surface of the antigen engaged B cell. Thus, if 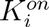 is the rate of association of antigen to B cell *B*_*i*_, and *1/τ* is the average rate at which a B-cell internalizes bound antigens, the ratio of peptide populations present on the surface of cells *B*_*i*_ and *B*_*j*_ is limited by the ratio of binding constants as:

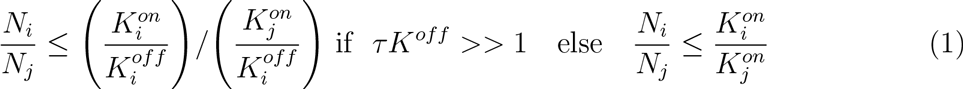

For perfect Tfh cell recognition and binding, Eq.(1) determines the optimal “equilibrium” rate at which B cells with different antigen affinities can be discriminated by Tfh cells, assuming similar efficiencies of antigen processing and epitope presentation. In principle, the ratio, Eq.(1), can be higher (by a maximum factor *p*) if the efficiency of epitope presentation is positively correlated with BCR-antigen binding affinity. Since in this scenario, B cells do not leave the LZ in the absence of a successful costimulation event, every B cell will eventually encounter enough Tfh cells until it obtains sufficient costimulation, the affinity discrimination ratio is unaffected by Tfh cell help and is given by Eq.(1). In addition, the mechanism, Eq.(1) does not discriminate between antigen-specific and cross-reactive B cells which recognize self antigens. Thus, a simple equilibrium model of B cell editing that depends solely on differential amounts of antigen binding and presentation is insufficient to satisfy the twin goals of high antigen affinity and discrimination against self antigen.

How can the limit, Eq.(1) be improved upon? We suggest that a natural mechanism of affinity discrimination would be to penalize B cells that take longer to obtain Tfh cell costimulation. In this context, it has been shown that a majority of the somatically hypermutated BCRs that reach the LZ undergo several encounters with Tfh cells, with only a few such encounters resulting in successful Tfh cell engagement.^3,13,14^ In addition, studies indicate that many B cells undergo apoptosis during the GC reaction,^3^ with some studies suggesting that B cell apoptosis is dependent on the amount of bound antigen and could serve as a mechanism for antigen discrimination.^15–17^

It is known that the extent of Tfh cell help depends on the level of antigen engagement,^9,18^ furthermore successful Tfh costimulation causes exit into the DZ from the LZ, and its level determines the subsequent extent of cell division and SHM.^5,18–21^ Thus, experimental evidence indicates that i) B cells appear to need a large number of B-Tfh cell reactions in order to form the right interaction with a Tfh cell for costimulation, ii) Antigen engaged and pMHC presenting B cells continually undergo apoptosis or migration from the GC light zone, and iii) there is a substantial fraction ( 15 — 30%) of B cells that return to the dark zone for further expansion.^3,5,13^

The experimental studies and the analysis of equilibrium discrimination that we have discussed suggest that affinity selection of antigen bound B cells in the light zone is due to a competition between the binding of B cells to Tfh cells and loss of B cells from the GC light zone, either due to apoptosis, or due to a return of B cells to the GC dark zone. We propose that this competition is the fundamental mechanism that underlies affinity selection of B cells.

### A. Antibody Repertoire Editing: The role of B cell loss

We present a simple analytical model to show that this competition is sufficient to substantially enhance affinity discrimination and also allow editing of cross-reactive B cells. The model we describe is summarized in Fig.1.

B cells with BCR sequence s that are activated upon binding to a number *N*_*a*_*(s)* of antigens on follicular dendritic cells (FDCs), on average display a number of peptide epitopes, *n*_*p*_*(s)*. For simplicity we ignore any intrinsic cell-cell variation in the number of peptide epitopes displayed, even for a given antigen affinity. We instead assume that the average number of epitopes, *n*_*p*_*(s)*, is representative of the actual epitope population displayed. We initially assume that the epitope population consists of a single peptide sequence, and this is discarded in the subsequent analysis. We assume that the average number of antigen epitopes displayed is a function of the amount of bound antigen, i.e *n*_*p*_(*s*) ≡ *n*_*p*_[*N*_*a*_(*s*)].^5,20,22–24^

Following successful antigen binding, B cells undergo chemotaxis in zones rich in Tfh cells. Depending on the type of GC, Tfh cells can either be localized at the GC light zone periphery or present in the GC light zone along the FDC’s. The intermediate states *i* ∈ {0,1, 2,…, *N* − 1} of an antigen bound B cell are most simply described as a count of the number of B-Tfh encounters that the cell has previously undergone. We summarize the B-Tfh cell encounter reaction by the scheme, Fig. 1(b), with rates that depend on the B
cell state:

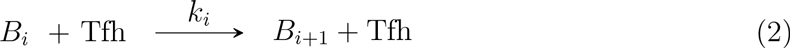

**FIG. 1.**
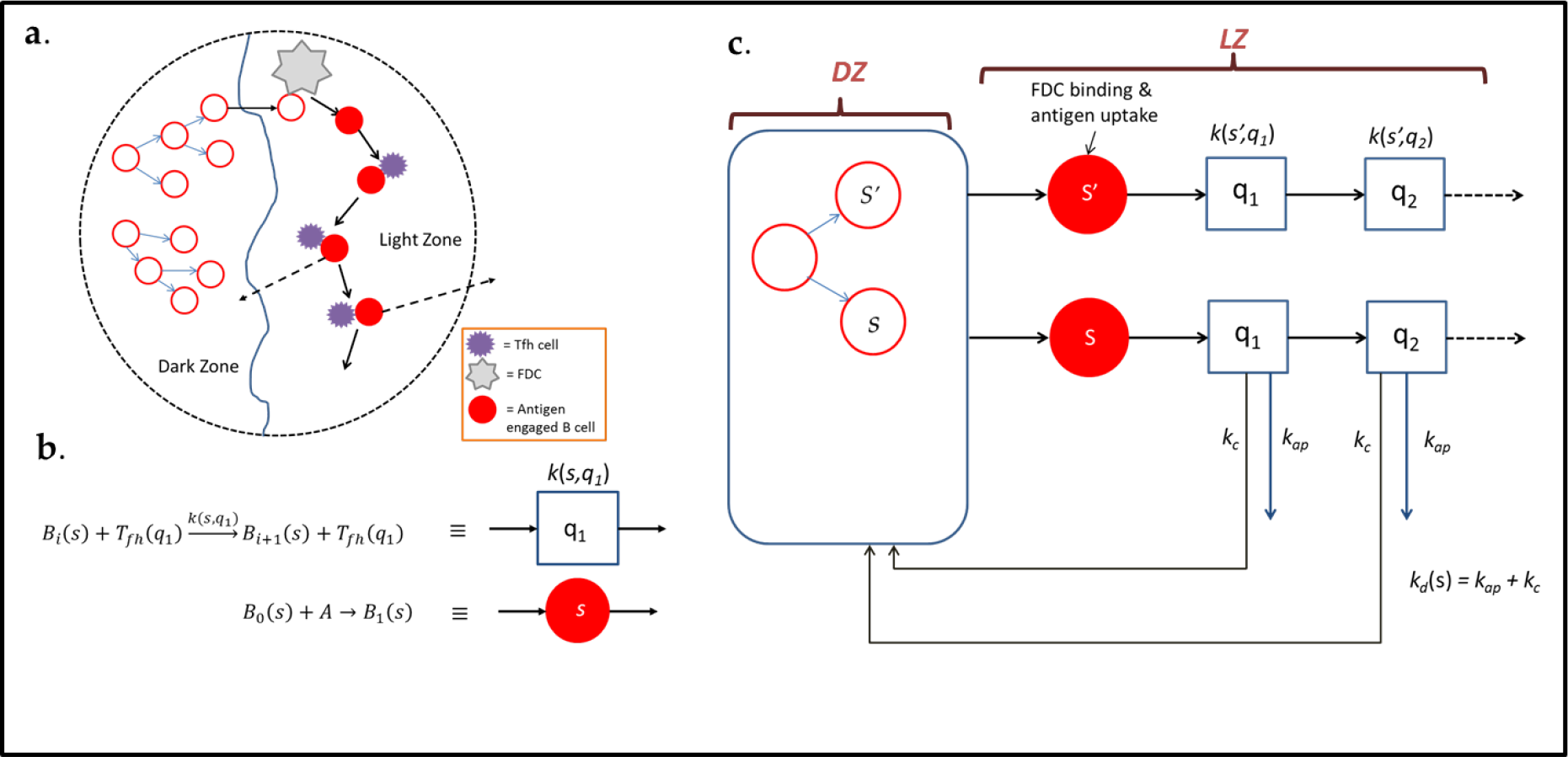
(a) A sketch of the GC B cell reaction. Open red circles are antigen-free B cells while filled circles are antigen engaged B cells. The arrows represent B cell division accompanied by SHM. (b) Schematic representations of individual B cell encounters with follicular DC’s and Tfh cells. (c) A pictorial description of successive B cell encounters and fate in the GC.

Here *k*_*i*_ ≡ *k*(*s*, *q*_*i*_) is the effective rate constant of the reaction that could generally depend on the concentration, *B*_*i*_, and sequences *s*, of B-cells, *q*_*i*_ of the cognate T-cell receptor, in state *i*. In addition B-cells in each state undergo apoptosis or exit from the GC light zone at a BCR sequence dependence rate, *k*_*d*_*(s)*. The rate, *k*_*d*_*(s)* is a sum of the apoptosis rate, *k*_*ap*_ and a rate of exit from the GC light zone, *k*_*c*_, i.e *k*_*d*_*(s)* ≡ *k*_*ap*_ + *k*_*c*_. The loss of B cells from the GC light zone reaction can be described by the scheme (Fig. 1(c)):

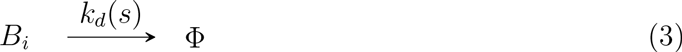

It is to be noted that the B-Tfh reaction involves chemotaxis of B cells towards Tfh cells along a chemokine gradient, and the rates used in this work are assumed to include its effects.^25–29^ Intuitively, the likelihood that a B cell survives the duration between successive Tfh cell encounters is given by the ratio of its rate of successful engagement by a Tfh cell to the total rate at which the B cell changes state. Thus, we define a transmission probability of converting states *B*_*i*_ → *B*_*i*+1_ as

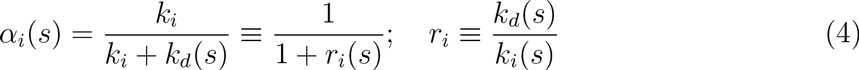

The population, *σ*(*B*_*n*_, *s*) of B cells that have undergone *n* encounters with Tfh cells at steady state is given by the product of its probabilities of surviving each of the *n* Tfh cell encounters:

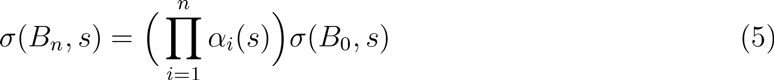

Thus, the ratio of two B-cell populations with BCR sequences {*s*, *s’*}, after *n* Tfh cell encounters is

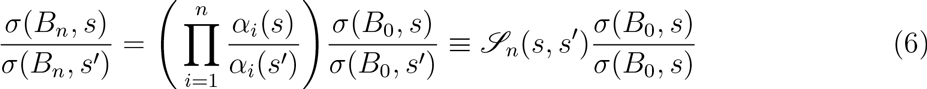

The selectivity, 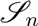 as described by Eq.(6) is the ratio of the population of B cells with BCR sequences {*s*, *s’*} after *n* encounters with Tfh cells assuming that their initial populations are equal. The ratio, *r*(*s*), measures the relative probability, and hence the competition between Tfh cell engagement and GC LZ B cell loss. As an example, for two BCR sequences such that *r*(*s*) = *ν* > 1 and 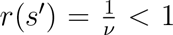, (i.e their rates of B cell loss are higher (lower) than rates of Tfh engagement), the selectivity becomes

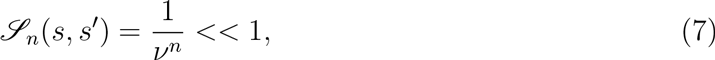

as the number of encounters, *n*, with Tfh cells increase. Thus, when B cells bind antigens with high avidity, they favor Tfh cell engagement over loss from the LZ, since they present more antigen, and also receive stronger anti-apoptotic stimuli. For such cells, we have *r*(*s*) ≪ 1, and vice versa when B cells have low avidity for antigen. In combination these two effects reduce the viability of B cells with BCRs of low affinity for selection. The differences between high affinity and low affinity BCR sequences are magnified exponentially through multiple encounters with Tfh cells prior to the determination of B cell fate, as shown by Eq.(6). From Eq.(6) the absence of B cell loss results in a selectivity ratio, 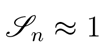 and multiple Tfh cell engagements do not result in any further discrimination between B-cells. This is consistent with our previous equilibrium analysis where for sufficiently long-lived B cells, Tfh cells do not confer much selectivity between different BCR sequences for their binding affinity. Hence, the loss of B cells from the GC light zone during affinity selection is necessary for achieving the enhanced selectivity in BCR affinity for antigen.

**FIG. 2.**
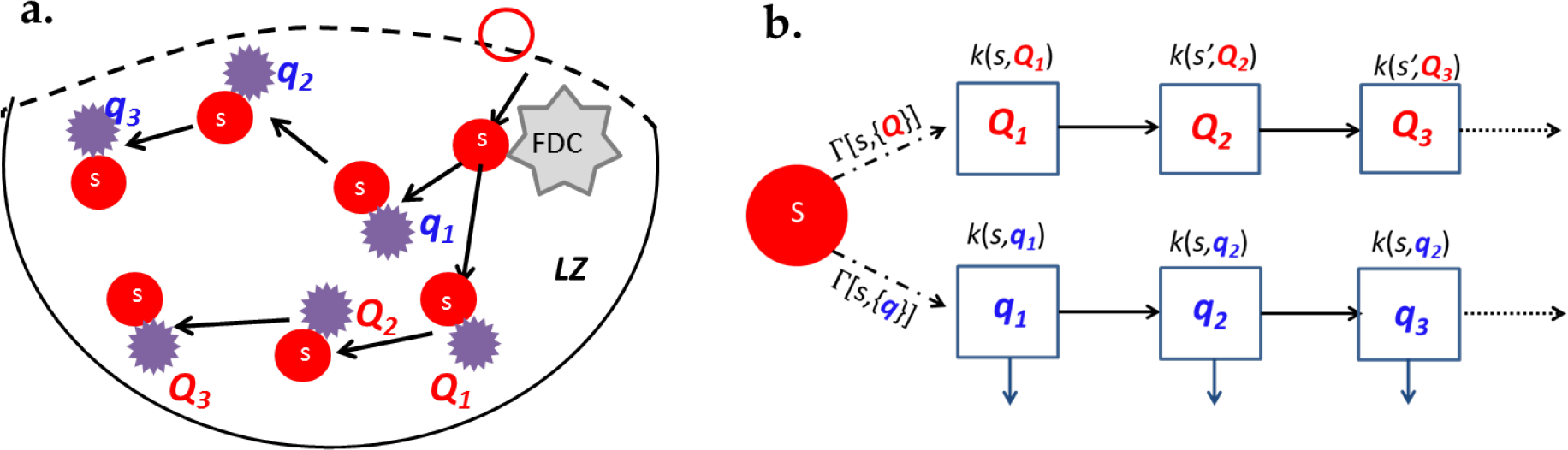
(a) A sketch of the possible pathways of B-Tfh cell encounters. Each B cell has a probability of encountering different sets of Tfh cells, e.g Tfh cells with sequences {*q*_1_, *q*_2_, *q*_3_,‥} or alternately encounters involving Tfh cell sequences {*Q*_1_, *Q*_2_, *Q*_3_,‥}. (b) Each such B-Tfh cell “path” can be represented by sequences of individual B-Tfh reactions as shown.

### B. Affinity Maturation with Variable Epitope Affinities and MHC Turnover

We have argued that when the rate of B-Tfh cell encounter is the limiting step in the GC light zone reaction that loss of B cells results in an exponential enhancement of BCR affinity selection. Remarkably, the model implies that this gain is realized even when the loss rate is independent of BCR sequence as long as the amount of antigen presented is positively correlated with BCR antigen binding affinity. However, these conclusions have been reached with the assumption that all the Tfh cells present in the GC light zone are from a single clone, and that the rates of B-Tfh cell encounter are identical. In general, the Tfh cell population in the GC is heterogenous in the TCR sequences present, although all the TCRs present can be assumed to recognize at least one of the non-self pMHC presented on the surface of GC B cells. We generalize our model of selection to account for these additional factors. The generalized dynamics are illustrated schematically in Fig. 2

Let 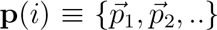 represent the pMHC epitope sequences that are presented on the surface of a GC B cell after *i* encounters with Tfh cells, 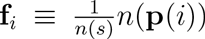 the vector of frequencies of each epitope presented on the B cell, *B_*i*_*, and of σ[**f**_*i*_;*n*(*s*)] represent the efficiency (or probability) of presenting pMHC complexes at frequencies **f**_i_ given a total number, *n*(*s*), of peptide epitopes. We assume that the efficiency of peptide presentation is independent of the number of peptides bound, i.e σ[**f**_*i*_;*n*(*s*)] ≡ σ[**f**_*i*_]. Similarly, we define the probability of a Tfh cell having its TCR sequence, *q*_*i*_, as *γ*(*q*_*i*_). From these definitions, the probability of a B cell with BCR sequence s surviving N encounters with Tfh cells is:

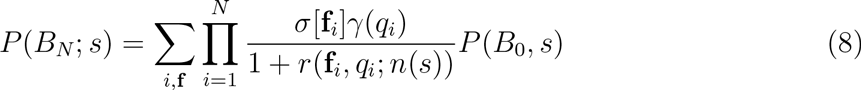

Here, as before, *r*(**f**_*i*_, *q*_*i*_;*n*(*s*)) is the ratio of apoptosis rate of a B cell, *B*_*i*_, to its rate of binding to a Tfh cell with TCR sequence *q*_*i*_. This ratio depends both on the sequences of pMHC complexes presented on the B cell surface and the specific TCR sequence presented on the Tfh cell. Furthermore, the rate of Tfh cell binding is proportional to the number of pMHC complexes that are complementary to its TCR. Thus,

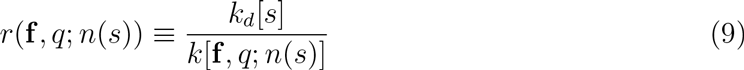

The binding constant *k*[**f**, *q*; *n*(*s*)] depends on the number of pMHC epitope complexes that can bind a Tfh with TCR *q*. Thus, a natural approximation to the binding constant is

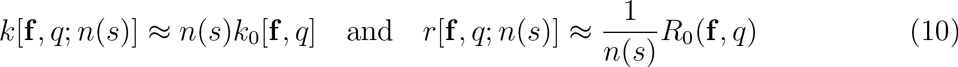

where the quantity, *k*_0_[**f**, *q*], in Eq.(10) depends only on the frequency of pMHC complexes present on the B cell surface, rather than their absolute numbers. The ratio, *R*_0_, has only a weak dependence on the number of bound antigens, which we henceforth ignore. Eq.(8) and Eq.(10) result in:

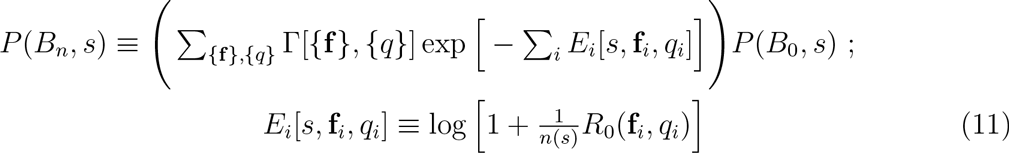

The function Γ[{**f**}, {*q*}] ≡ **Π**_*i*_ σ[**f**_*i*_]*γ*(*q*_*i*_), in Eq.(11) is the probability of *n* successive B-Tfh interactions with the *i*-th interaction being between an epitope distribution, **f**_*i*_, and a Tfh cell TCR sequence *q*_*i*_ assuming that each such interaction is always successful. As expected, when either the intrinsic encounter rate of B cells to Tfh cells is very high relative to their apoptosis rate, or B cells bind large amounts of antigen, Eq.(11) shows that there is an exponential enrichment of B cell survival probability for B cells that bind more antigen. To see this, we rewrite Eq.(11) using:

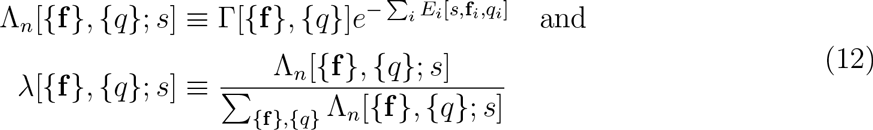

λ is the probability of a B cell with BCR sequence s executing a particular trajectory of *n* B-Tfh cell reactions. The ratio of survival probabilities in terms of λ is:

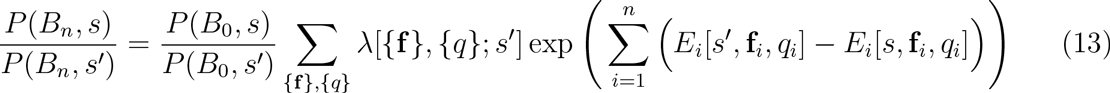

On defining Δ*E*_*i*_(*s*, *s’*) ≡ *E*_*i*_[*s*, **f**_*i*_, *q*_*i*_] − *E*_*i*_[*s’*, **f**_*i*_, *q*_*i*_], the relative probability is written in a physically transparent form as:

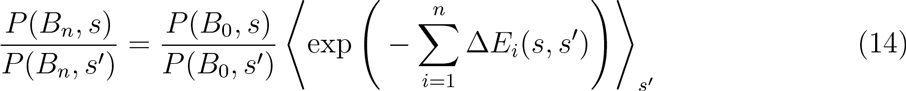

Here, ⟨..⟩_*s’*_ is the weighted average over the probability distribution, λ[{f}, {q}; *s’*], of a B cell with BCR sequence *s’* having *n* encounters with Tfh cells having all the possible kinds of TCR sequences {*q*}. When the amount of antigen bound to BCRs of sequence s is large, i.e when *n*(*s*) > *n*(*s’*), we have that Δ*E*_*i*_(*s*, *s’*) < 0 for every possible B-Tfh cell encounter. Thus, as the number of B cell encounters with Tfh cells increases, the relative survival probability of B cells with higher amounts of antigen increases exponentially. A numerical calculation of this selection process is shown in Fig. 3, which illustrates the degree of discrimination between B-cells with different BCR sequences, as a function of the number of Tfh cell encounters, while treating the ratio *r* as a random variable. We performed such calculations over a multiple range of values of the mean and variance, and also for different choices of the probability distribution, and the results remain robust to these choices (Appendix A).

**Fig. 3.**
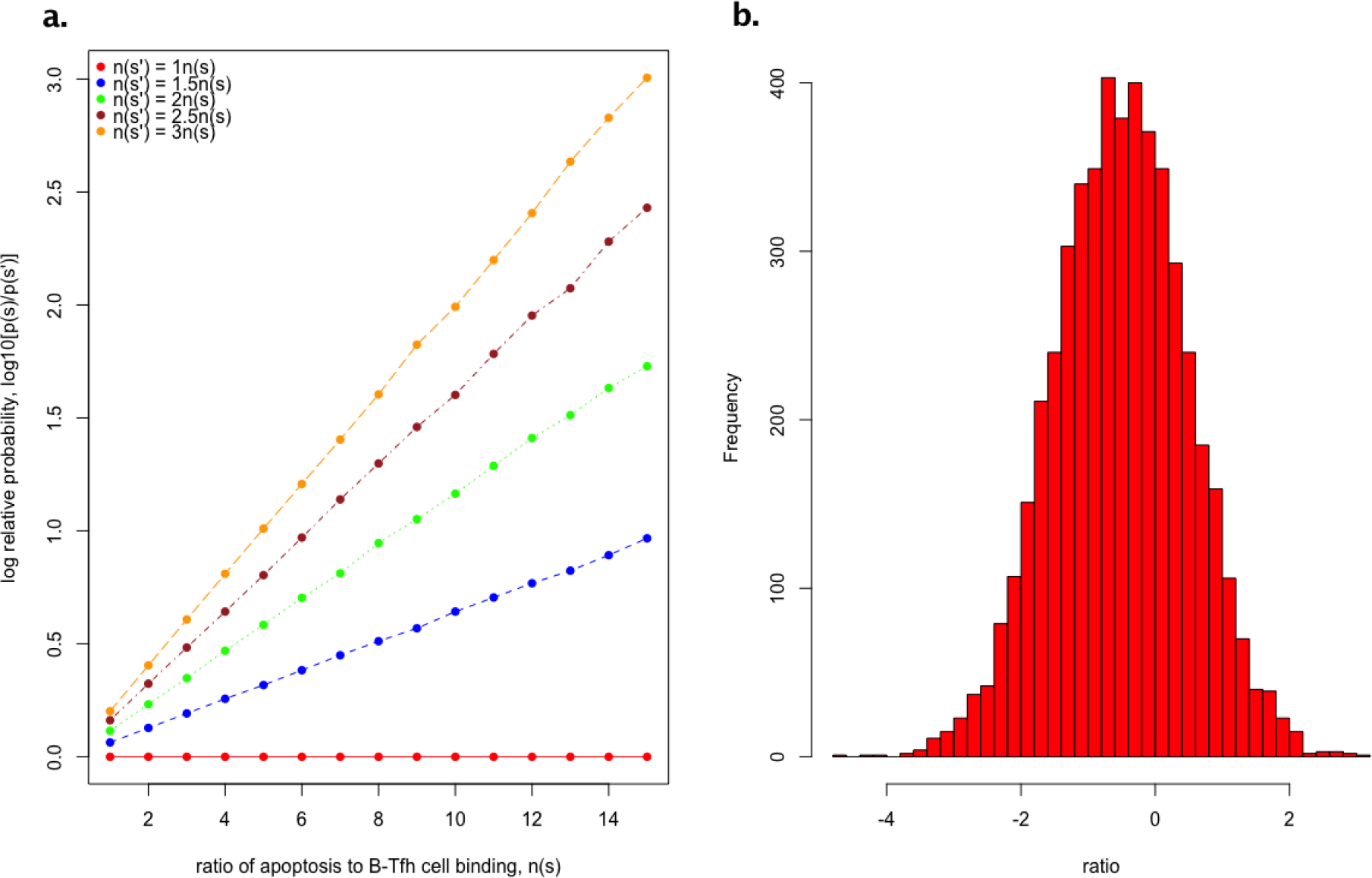
a). Logplots of the relative probability of B cells with BCR sequences *s’* and *s* as a function of the number, *n*, of B-Tfh cell encounters. To model individual B-Tfh reaction and apoptosis rates, we randomly sample the ratio of B cell loss rate to B-Tfh cell encounter rate constants, *r*[**f**, *q*; *n*(*s*)] from a lognormal probability distribution, with a mean −0.5 (i.e the rate of B cell loss is on average a third smaller than rates of B-Tfh cell engagement), and a variance of 1.0 to allow for the possibility of a range of B-Tfh cell encounters, as shown in panel b). About 5000 such trajectories are generated and the total probability obtained by summing over all such trajectories.

### C. Discriminating Self from Non-self

Our analysis argues that positive selection of B cells for antigen affinity occurs as a consequence of two factors. First, the quantity of pMHC presented on B cells is a function of the amount of bound antigen, with the amount of stable pMHC increasing with quantity of bound antigen, and second the accessibility of MHC presenting B cells to Tfh cell costimulation is a consequence of tradeoffs between binding to Tfh cells and B cell apoptosis. Here, we show that the same basic mechanism could also discriminate against self reactive B cells. Fig.4 illustrates two scenarios where cross reactivity could affect B cell selection in the light zone. In the first scenario (Fig. 4(a)), there are cross reactive B cells that present both self and non-self antigens in the absence of any self reactive Tfh cells, while in the second scenario (Fig. 4(b)), such B cells are also recognized by self reactive Tfh cells. We examine the role of B cell loss in maintaining B cell tolerance in both these scenarios.

**Fig. 4.**
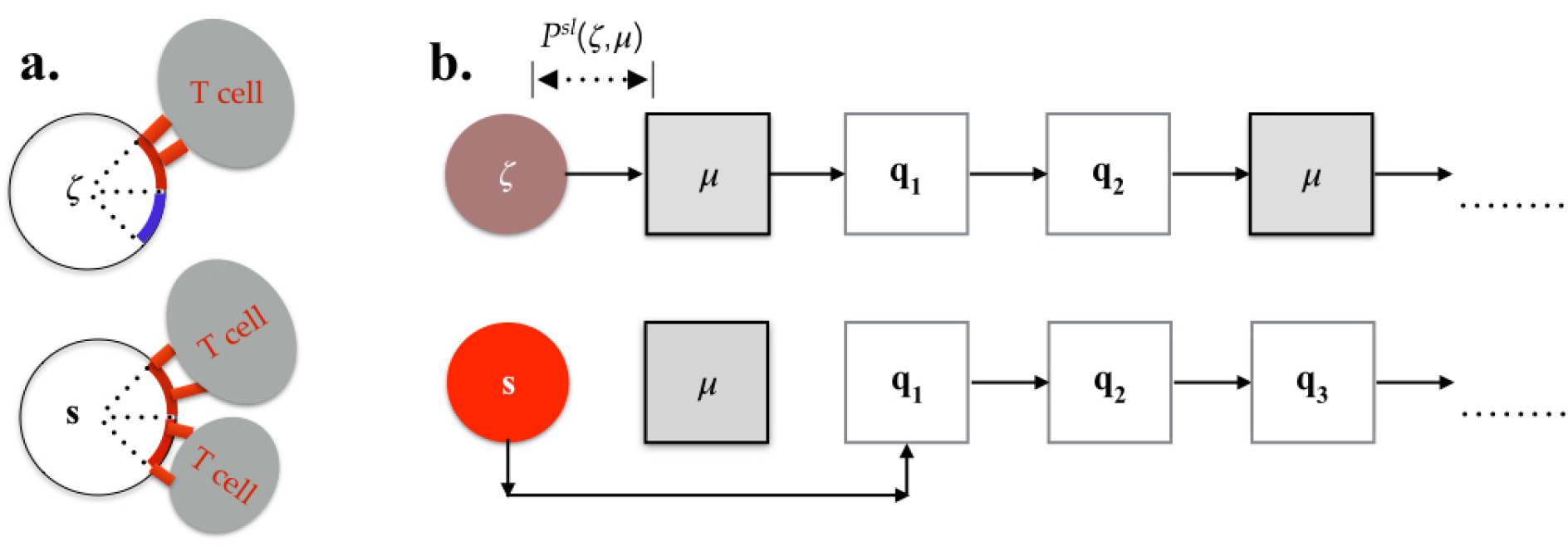
(a) A sketch showing an autoreactive B cell with half of its pMHC consisting of self antigens (the blue arc) and the other half consisting of non-self pMHC (red arc). Tfh cells (ellipses with TCRs as red rectangles) recognize only the non-self pMHC. (b) Cross reactive B cells can recognize autoreactive Tfh cells (grey boxes) while antigen specific B cells don’t, leading to different survival probabilities. A particular sequence of non-self and self Tfh cells that react with the B cells is shown as an example.

#### 1. *Tolerance to cross reactive B cells*

Consider B cells with BCR sequence *s* that have specific and high antigen binding affinity, and a second class of B cells with BCR sequence *ζ* that are cross-reactive with antigen and an arbitrary set of self antigens, as illustrated by Fig. 4(a). Let {*n*(*s*), *n*(*ζ*)} represent the total amount of bound antigen, either non-self or self, and **F** ≡ {**f**_*self*_, **f**_*ns*_} represent the frequency vector of self (**f**_*self*_) and non-self (**f**_*ns*_) pMHC’s presented on B cell with BCR ζ. From Eq.(10) the binding constant of the cross-reactive B cells to a Tfh cell with TCR sequence *q* is

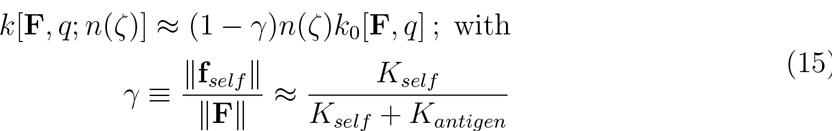

The norm, ‖**F**‖, in Eq.(15) is the total number of pMHC complexes presented per quantity of bound antigen (self or non-self), and ‖**f**_*self*_‖ is the total fraction of self antigen pMHC complexes presented. Thus, if the total amount of bound antigen is the same, i.e if *n*(*s*) ≈ *n*(*ζ*), the proportion of antigenic pMHC presented on the surface is smaller by a factor (1 − γ) and from Eqs.(13) and (14), the ratio of survival probabilities after m B-Tfh encounters is:

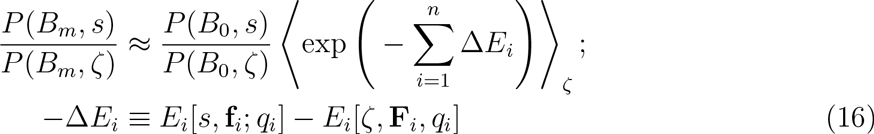

The “energies” in Eq.(16) satisfy 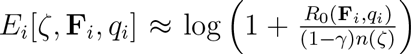. Since cross-reactive B cells can display both self and non-self pMHC, they have a different peptide presentation profile from antigen-specific B cells. At the least, the fraction of antigenic pMHC’s presented is reduced because of the additional presence of self pMHC’s. Thus in general, *R*_0_(**F**_*i*_, *q*_*i*_) ≤ *R*_0_(**f**_*i*_, *q*_*i*_) and (1 − γ)*n*(ζ) < *n*(*s*) resulting in the energy difference Δ*E*_*i*_[*s*, *γ*] < 0 for every encounter *i*. This results in an enrichment of antigen specific B cells with sequence *s* in comparison to cross-reactive B cells with sequence ζ as shown by Eq.(16).

When cross-reactive B cells bind to self antigen with greater or comparable affinity to the foreign antigen, the factor γ increases and the effective amount of foreign antigen pMHC presented becomes small, and Tfh costimulation harder to obtain. Consequently, Tfh cell costimulation will result in selection against the cross-reactive B cell and in favor of foreign antigen specific B cells with comparable, or higher affinity for foreign antigen. On the other hand, B cells with high affinity for foreign antigen but low affinity for self antigen would still be favorably selected by Tfh cell costimulation. This is consistent with experimental observations, wherein the affinity matured population of B cells include those that have weak cross-reactivity towards self antigens. This analysis suggests that the negative selection of cross-reactive B cells is possible only if self-antigens are efficiently presented for possible uptake by B cells, so that they can compete with the uptake of foreign antigen in the GC. One mechanism for efficient self antigen presentation is to concentrate self-antigens in localized regions of the GC, as observed in recent experiments.^12,30^ In the absence of efficient self-antigen presentation, GC affinity selection will be unable to distinguish between crossreactive and antigen-specific B cells.

#### 2. *Tolerance and imperfect Tfh cell repertoire editing*

Peripheral tolerance in the GC can also be broken by Tfh cells that recognize self pMHC complexes due to imperfect Tfh repertoire editing (Fig 4(b)). Here we examine the robustness of B cell affinity maturation mechanisms in the presence of self recognizing Tfh cells. For illustrative purposes consider that only a single self recognizing Tfh cell with TCR sequence *μ* is present in the GC light zone,at a fraction *δ* of the total number of Tfh cells present in the GC light zone. We assume that the rate of encounter between the self reactive Tfh cells and B cells with sequence *ζ* is constant and independent of the number of Tfh cell encounters, For example a cross-reactive B cell with BCR sequence *ζ* has a probability of having *ν* Tfh cell encounters such that two of these are autoreactive T cells (Eqs.(B1)-(B3)):

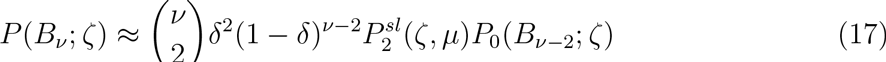

Here, *P*_0_(*B*_*ν-k*_, *ζ*) is the probability of encountering ν — *k* antigen specific Tfh cells among all the possible combinations in which *n* — *k* B-Tfh cell encounters can occur, in the absence of any self reactive Tfh cells in the GC, and 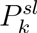 is the probability of *k* encounters between the self recognizing Tfh cells and the B cell. In addition, if we assume that individual B-Tfh cell encounters are independent events, we can approximate *P*_0_(*B*_*n*_, *ζ*) ≈ *P*_0_(*B*_*k*_;*ζ*)*P*_0_(*B*_*ν-k*_;*ζ*). Using this approximation, the ratio of probabilities of *n* B-Tfh encounters between a cross reactive B cell in the presence and absence of self reactive Tfh cells is:

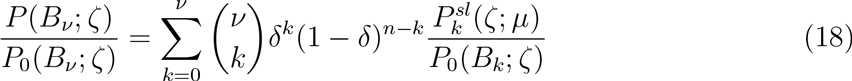

Since we have assumed that there is only one type of self recognizing Tfh cell present, the probability is a product of identical probabilities of single encounters with self Tfh cells, i.e 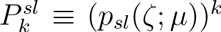 If *p*_0_(*ζ*) and *p*_*m*_(*ζ*) are lower and upper bounds respectively on the effective probability of a single B-Tfh cell interaction in the absence of any self reactive Tfh cells, we can show that (Eq. (A9)):

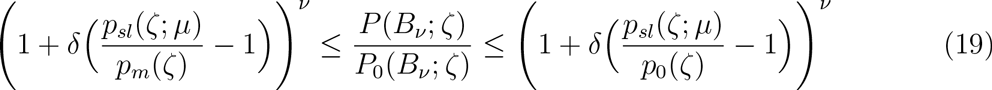

The inequality at the right hand side of Eq.(19) shows that when the probability of binding to a self Tfh cell is higher than the smallest probability of binding to a non-self Tfh cell, self Tfh cells could successfully compete with antigen specific Tfh cells to provide costimulatory signals to cross reactive B cells, and correspondingly antigen specific B cells are favored when the probability, *p*_*sl*_, is less than the highest effective probabilty of encounter with antigen specific Tfh cells.

The ratio of survival probabilities of B cells that are cross-reactive against self antigens, and those that are antigen specific B cells can be estimated from Eqs.(14) and (19). Let *s* represent the BCR sequence of an antigen specific B cell. Then, we write:

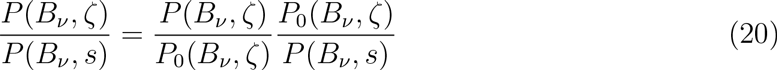

The physical interpretation of *P*_0_(*B*_*ν*_, *ζ*) is that it is the probability of *ν* encounters with only antigen specific Tfh cells. From Eqs.(14) and Eqs.(19)-(20), we obtain the following expression:

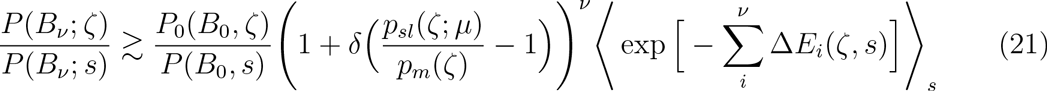

An inequality in the opposite direction as Eq.(21) holds when *p*_*m*_ is replaced by *p*_*0*_. Eq.(21) is recast upon defining average “energy” differences over each sequence of B-Tfh cell encounters as:

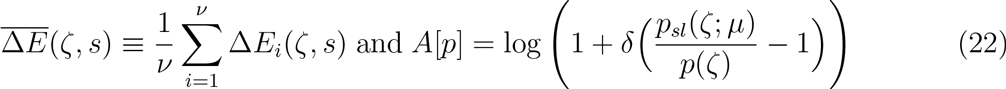

We rewrite the inequality, Eq.(21) as

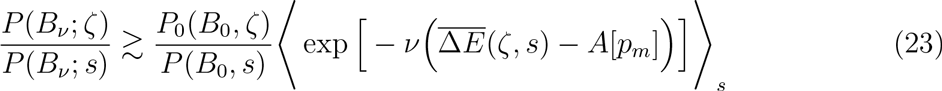

Since the amount of self antigen presented by the B cell ζ is γ*n*(*ζ*), from the definitions,
Eq.(8):

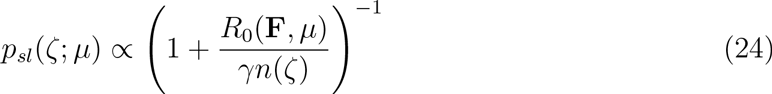

and *p*_*m*_(*ζ*) has a corresponding dependence on (1 − γ)*n*(*ζ*). It can be seen from Eqs.(23) and (24) that when the amount of self antigen presented is the dominant proportion of the total amount of antigen presented by the B cell, *p*_*sl*_ > *p*_*m*_and *γ* ≲ 1. In this situation, cross reactive B cells are preferentially selected over those B cells which present even smaller amounts of non-self antigen, i.e (1 − γ)*n*(*ζ*) ≥ *n*(*s*). This means that tolerance is preserved unless the cross reactive B cells present both a high amount of non-self antigen and a substantial greater quantity of self antigens, i.e they have very high binding affinity for both self and non-self antigens. However, cross-reactive B cells can compete for selection with antigen specific B cells, when roughly equivalent amounts of self and non-self antigen are presented by the cross-reactive B cells such that *p*_*sl*_ ≈ *p*_*m*_and *n*(*s*) ≈ *n*(*ζ*)(1 − *γ*). In either case, by the uptake of self antigen, cross reactive B cells are penalized in their encounters with antigen specific Tfh cells due to the presentation of a reduced number of non-self antigens on their surface. This penalty can only be partially compensated for by the presence of self reactive Tfh cells, since their overall ability to obtain enough costimulation to leave the light zone would require simultaneous stimulation from, and polarization of, both self and non-self Tfh cells.

## III. EFFECTS OF LIGHT ZONE SELECTION ON B CELL CLONAL EXPANSION IN THE GC

We have described a mechanism by which selection for high affinity BCRs can occur in the GC LZ. We now consider the effects of such a selection mechanism on the clonal diversity generated in the GC DZ during B cell division and mutation. B cell clonal expansion in the GC DZ occurs through asymmetric cell division wherein BCRs in one of two daughter cells acquire a mutation at each division. From Eqs.(11)–(12), given an initial number of B cells, *N*(*B*_0_, *s*) with BCR sequence, *s*, that are yet to encounter Tfh cells in the GC LZ, the number of B cells that remain in the LZ after ν Tfh cell encounters is:

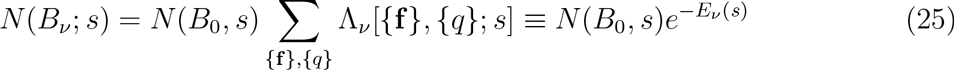

We assume that the probability, *ρ*(*ν*, *s*), of returning to the DZ from the LZ is a function of their number of Tfh cell encounters and level of costimulation. We can treat the clonal expansion of B cells in the GC dark zone as a branching process, and estimate the average number of clones upon clonal expansion. Thus, the number of clones of *N*(*B*_*ν*_, *s*) B cells that enter the dark zone is, (Eq. (C5)):

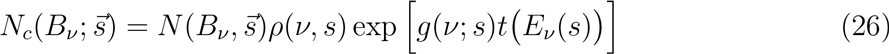

Here, we have assumed that the effective (average) growth rate, 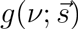 of all B cells that are descended from an initial B cell with BCR sequence *s*, depends on the number of Tfh cell encounters of the ancestor cells (their “costimulatory state”). The average residence time, in the GC dark zone, of B cells after *ν* Tfh cell encounters is 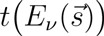, which depends on the extent of Tfh cell costimulation,^9^) 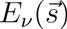. Thus, the number of B cell daughter cells from a parent BCR sequence, 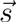 is

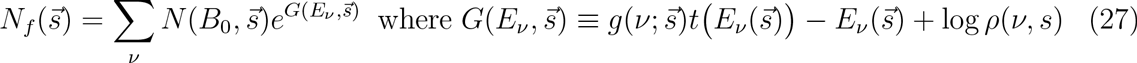

Eq. (27) shows that there is a competition between the amount of Tfh cell costimulation that a B cell receives, and its ability to divide once it returns to the GC dark zone. The probability that a B cell undergoes multiple Tfh cell encounters reduces exponentially as the number of such encounters increase, while its growth rate upon returning to the dark zone increases correspondingly. For B cells that return to the dark zone after only a few Tfh cell encounters, they are more likely to have received insufficient costimulation, and correspondingly their growth rate is also limited. On the other hand, B cells that have many Tfh cell encounters are likely to have higher levels of Tfh cell costimulation, and thus an elevated growth rate, with a penalty that the probability of surviving multiple such encounters is exponentially small. Thus, the dominant contribution to Eq.(27) is from B cells that undergo a number of encounters *ν*\* that maximizes the exponent, i.e:

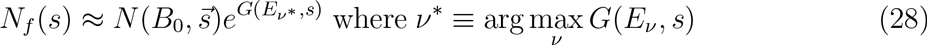

Under this approximation, the probability of clones originating from a sequence 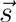 is given by

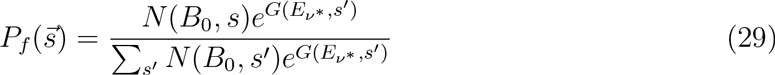

The optimal number, *ν*\* ≡ *ν*\*(*s*) of Tfh cell encounters for a B cell with a given BCR sequence, s, is a function of the number of pMHC complexes expressed on the surface of the B cell, i.e *ν*\* = *ν*\* [*n*(*s*)], since B cells that express higher amount of pMHC complexes can obtain costimulation after fewer Tfh cell encounters. Thus, we have that

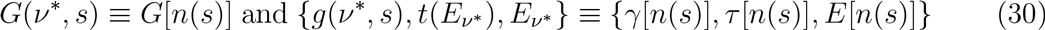

The growth rate and average duration of B cells in the dark zone are both increasing functions of the number, *n*(*s*) of pMHC complexes on the surface of the B cell. Similarly, B cells which have higher numbers of peptide epitopes presented survive longer and find it easier to get costimulation from Tfh cells. Thus, the optimal average duration, survival probability and growth rate satisfy the following conditions:

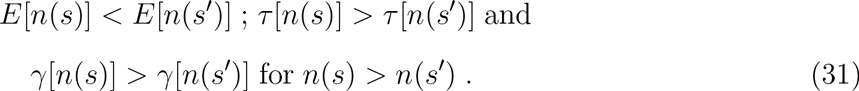

These properties collectively imply that for *n*(*s*) > *n*(*s’*), *G*[*n*(*s*)] > *G*[*n*(*s’*)] and consequently *P*_*f*_(*s*) > *P*_*f*_(*s’*). In particular, the difference in probabilities can be written in terms of their exponents:

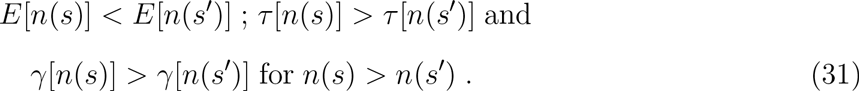

Let *s*^†^ be a BCR sequence which presents the highest amount of pMHC, i.e *n*(*s*^†^) = max_*s*_ *n*(*s*). We can then rewrite the probability of clones originating from a sequence *s* as

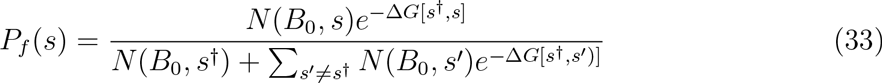

For *s* near *s*^†^, the exponents can be Taylor-expanded, such that with 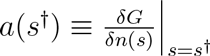, the probability of a B cell clone originating from such BCR sequences is:

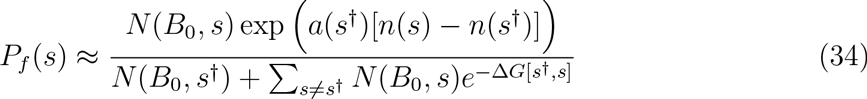

It is clear from Eqs.(33) and (34) that the probabilities of clones originating from sequences other than B cells with the highest antigen binding affinity and presentation efficiency are exponentially reduced. Thus, passage and clonal expansion of B cells through the DZ after selection by Tfh cells result in a further amplification of B cell clones that originate from those B cells that present maximal amounts of antigen pMHC complexes, and indirectly the highest antigen binding affinity. If the initial distribution of B cells include BCR sequences that have varied differences in antigen binding affinity, clonal expansion in the DZ amplifies this difference, through exponential increase in the number of clones originating from the higher affinity BCR relative to those originating from low affinity sequences. If there is a single BCR with substantially higher affinity than the others, then clones originating from such B cells will dominate, while heterogenous clonal distributions are obtained if there is more than one high affinity BCR sequence in the cells entering the DZ. This is consistent with studies by Tas et.al^31^ that demonstrate both heterogeneity in GC’s, with clones originating from multiple BCR sequences, or clonal domination from a single BCR sequence.

Our analysis of clonal expansion in the GC DZ is based on the average number of clones produced after multiple divisions. However, both the traversal of the GC LZ and subsequent entry into the DZ for clonal expansion are stochastic processes, resulting in a random number of daughter clones at the end of each generation time, whose variance increases with the number of generations. This stochasticity in the clonal expansion process also contributes to the heterogeneity of the clonal distribution in the germinal center, and allows for a finite probability of imperfect clonal amplification wherein clones from high affinity B cells are not the dominant population after clonal expansion. A fuller analysis of these possibilities is deferred here, since this is beyond the scope of this work and would need detailed numerical modeling.^32^ However, the elementary analysis presented here is sufficient to illustrate that the selection mechanism of B cells for their affinity by Tfh cells in the light zone proposed here is sufficient to explain the mechanisms of affinity selection in the light zone, and the observed patterns^31^ of clonal expansion in the dark zone.

## IV. DISCUSSION

We have proposed using very general arguments that selection of B cells in the germinal center by Tfh cells occurs due to a competition between the processes of Tfh cell recognition and B cell apoptosis/exit from the GC light zone, where B cells that present more antigen are ability to survive longer in the LZ, and also have an increased chance of a successful costimulation. Thus, affinity discrimination between B cells is predicted to be indirect, by selection in favor of B cells that present more antigen epitopes to Tfh cells. Recent studies have shown that indeed, Tfh cell binding and costimulation depends on the amount of pMHC complexes presented by cognate B cells^2,21^. However, due to this indirect discrimination, B cells that bind lower amounts of antigen but whose pMHC epitope presentation efficiency is high enough to compensate for reduced binding, are also predicted to undergo positive selection. This is consistent with experiments which demonstrate that Tfh cell selection of B cells depends only indirectly on antigen affinity, through the amount of pMHC presented on B cells.^3–5,9^ We note that our proposed selection mechanism has aspects that are similar to kinetic discrimination models used in other areas of biology.^33–35^

While B cells are selected for enhanced antigen presentation, and thus indirectly, antigen affinity, by Tfh cells in the LZ, the population differences due to this selection process are predicted to be amplified upon clonal expansion in the GC LZ. Our model predicts that clones of B cells with high antigen presentation are preferentially expanded due to a combination of effects, wherein greater presentation enhances both their survival and ability to be costimulated in the LZ, and a concomitantly greater duration, and number of cell divisions in the DZ.

We have argued that cross-reactive B cells undergo negative selection in comparison to purely antigen specific B cells because, at the very least, such B cells are able to express lower numbers of antigen-specific pMHC complexes in comparison to more antigen-specific B cells. However, the analysis predicts that B cells which bind large amounts of antigen but are weakly self reactive will be positively selected, implying that affinity selection against self reactive B cells selects for B cells that bind antigen more strongly than they bind any self ligands. The presence of self recognizing Tfh cells can also enhance the selection of cross-reactive B cells if present in numbers sufficient to offset the negative selection of such cells by apoptosis. This work therefore suggests that B cell affinity maturation occurs in a multistage process, with Tfh cells selecting for B cells with high antigen affinity as well as an ability to present antigen peptides efficiently, while the following clonal expansion in the DZ amplifies differences between selected B cell populations according to their affinity for antigen.

## ACKNOWLEDGMENTS

We thank Dr. S. Somani and Dr. G. Rajagopal for critical comments on the manuscript.

## COMPETING INTERESTS

V.K and K.E.B are both employed by and own stock in Janssen Research and Development LLC. which makes drugs to treat immune diseases. However, this work has no commercial relevance.

## APPENDIX A: Simulations of Discrimination between B cells

We tested the effects on the discrimination between B cells of sampling the ratio of B cell loss rates to B-Tfh cell engagement from different probability distributions, in addition to the lognormal distribution used in the main text. We found that the choice of this probability distribution does not alter the property of B cells to be discriminated according to the amount of antigen presented. As an example, we show in Fig.(5), the relative probability between two B cells as a function of number of encounters, for the ratio of B cell loss to B-Tfh engagement sampled from a beta distribution. We also examined whether discrimi-nation between two B cells with BCRs *s* and *s’* improves as the relative amount of antigen 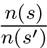 increases. This is plotted in Fig.(6), for different numbers of B-Tfh encounters. We can see that the relative probabilities increase by upto 4 orders of magnitude as 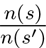 increases from 1 to 10. The increase in relative probability is also faster for B cells that undergo more as the mean ratio of apoptosis to B-Tfh engagement rates is altered. We examined this behavior for a ratio of antigens presented, 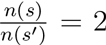, while maintaining the variance of the lognormal distribution from which the ratio of B cell loss rate to Tfh engagement rate is sampled from to be unity. We can see from Fig.(7) that the gain in relative probabilities plateaus as the rate of B cell loss increases relative to the rate of B-Tfh cell engagement. This is to be expected, since from Eq.(14) as this ratio increases, the relative probabilities of B cells of sequences s and s’ become independent of the relative rates of B cell loss to Tfh cell engagement, and rather depend mostly on the relative amounts of antigen presented and the number of B-Tfh cell encounters. Indeed, this can be seen in Fig.(7), where the relative probabilities plateau at lower levels for fewer B-Tfh cell encounters. We also have provided a R script from which these graphs were generated, for any further analysis of interest.

**Fig. 5.**
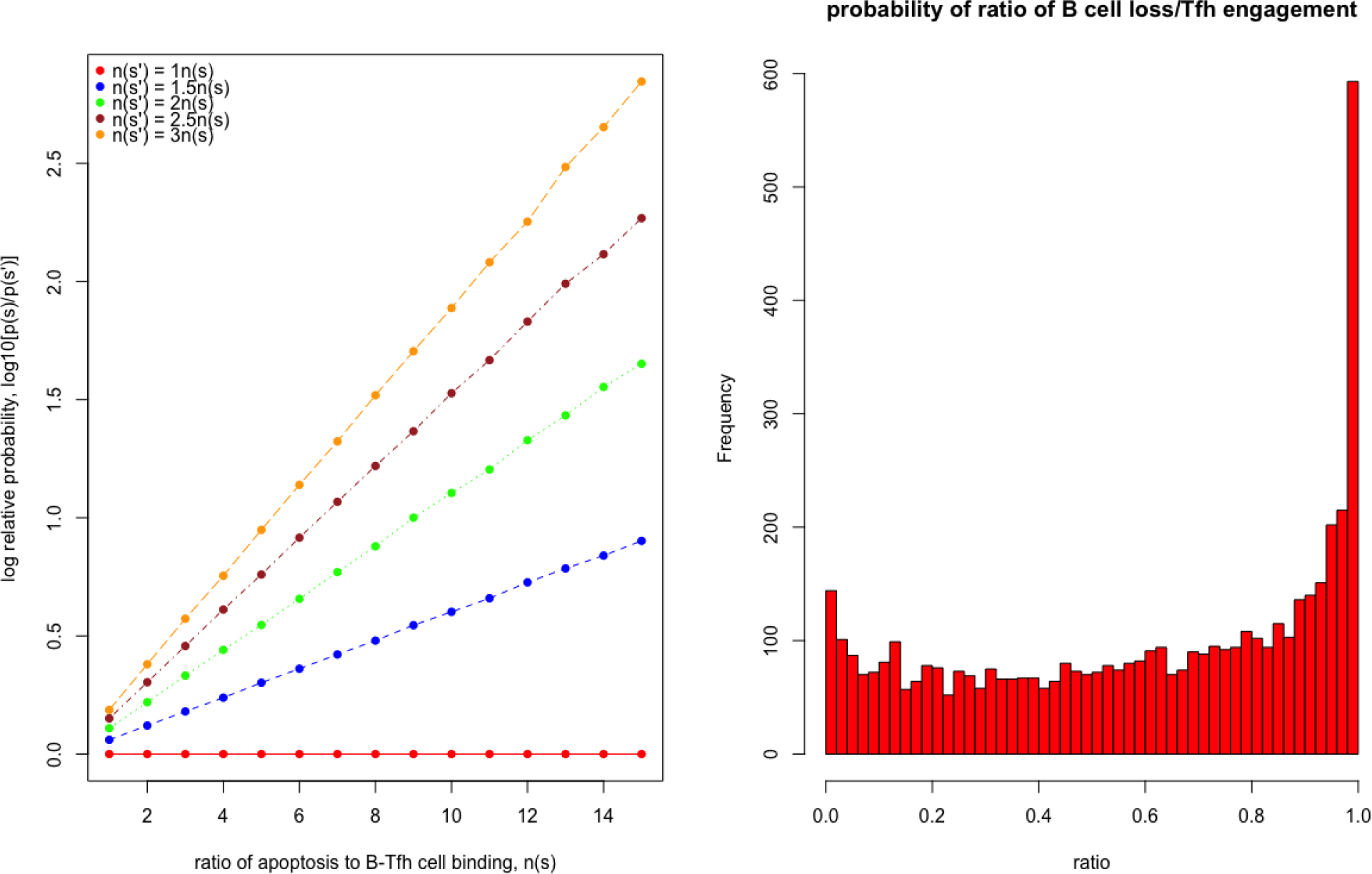
A graph showing how the relative probabilities of two B cells change as the number of B-Tfh cell encounters (a), where the ratio of B cell loss to Tfh cell engagement is sampled from a beta distribution shown in panel (b).

**Fig. 6.**
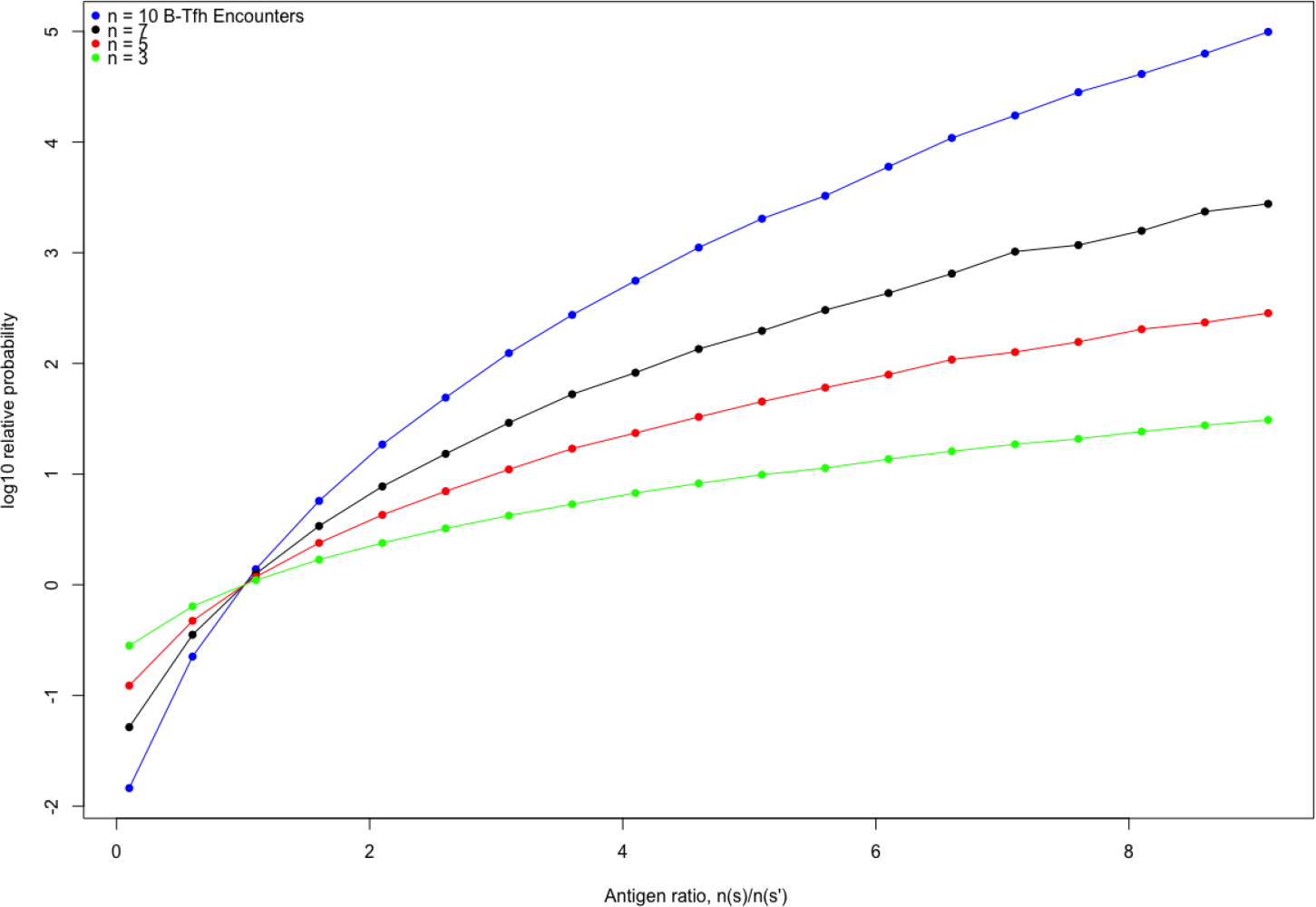
A graph showing how the relative probabilities of two B cells change as the relative amounts of antigen presented by the two cells change. The ratio of B cell loss to Tfh cell engagement is sampled from a log normal distribution with mean of −0.5 and variance of 1.0 as before.

**Fig. 7.**
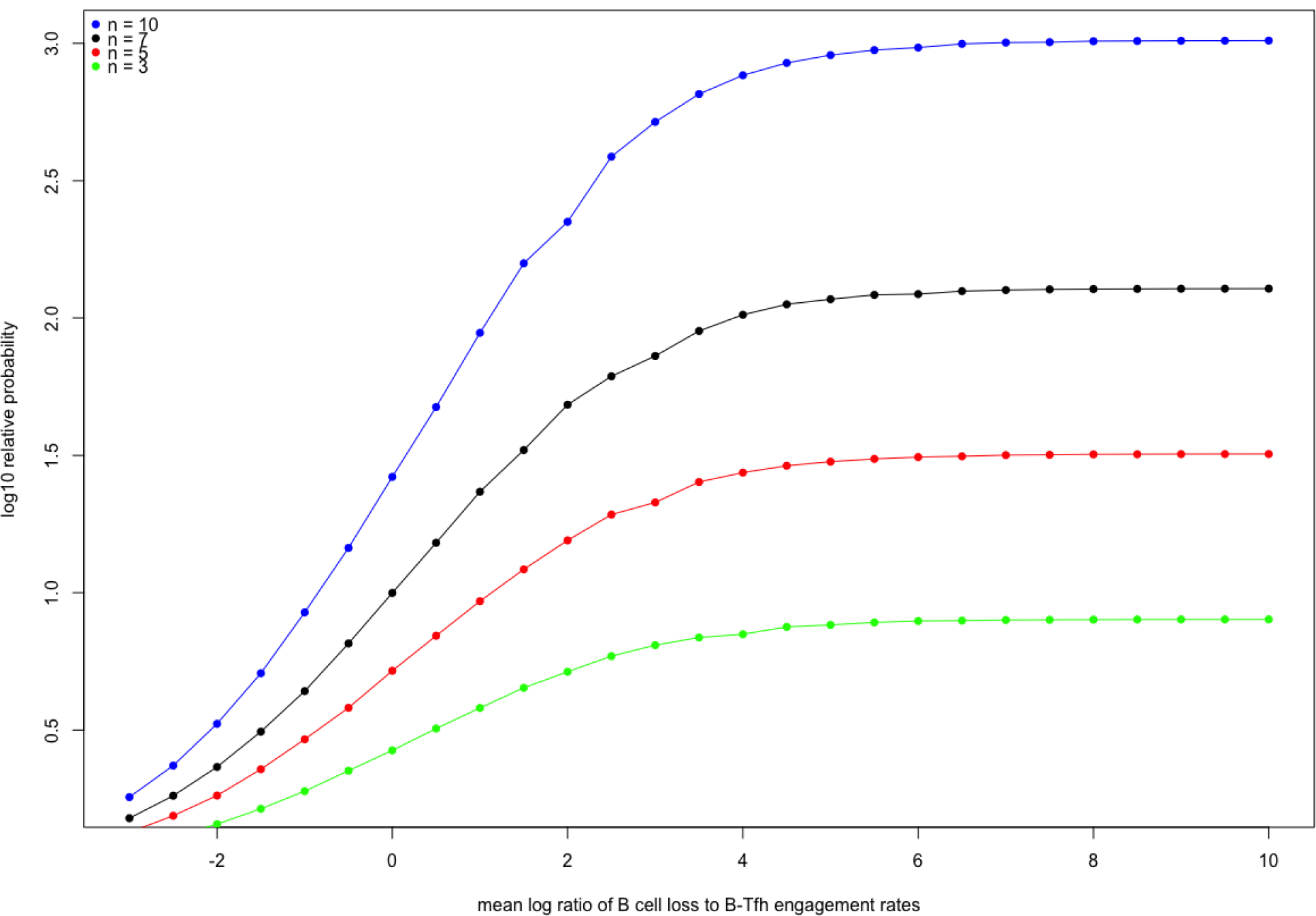
Relative probability of survival in units of *log*10 as a function of the mean value of log(*r*[**f**, *q*; *n*(*s*)]), the log of the ratio B cell loss to Tfh cell engagement. This ratio is sampled from the log normal distribution with the mean value plotted on the x-axis, and variance 1.

## APPENDIX B: Tolerance to imperfect Tfh cell repertoire editing

Let μ denote the TCR sequence of self reactive Tfh cells,at a fraction δ of the total number of Tfh cells present in the GC light zone. We can show that the rate of encounter between the self reactive Tfh cells and B cells with sequence ζ is constant and independent of the number of Tfh cell encounters, and that the probability of encountering *k* arbitrary antigen specific Tfh cells is the same for all choices of *k* Tfh cells. A cross-reactive B cell with BCR sequence ζ has a probability of having *m* Tfh cell encounters:

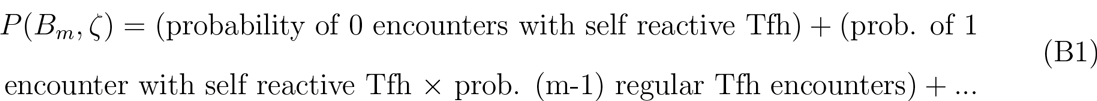

The probability of a single self-reactive Tfh cell encounter and *m* − 1 regular Tfh cell encounters is:

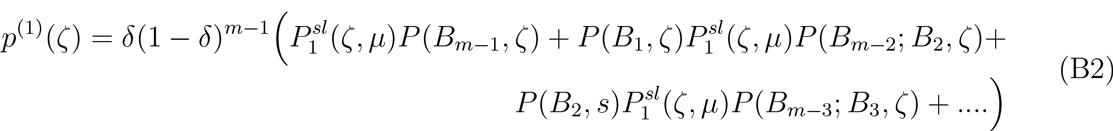

Here the probabilities *P*(*B*_*m-k*_; *B*_*k*_, *ζ*) are the probabilities of a B cell that has been previously activated by *k* encounters with Tfh cells undergoing *m* − *k* encounters with regular Tfh cells. If we assume that each B-Tfh cell encounter is independent of any that occurred earlier, we can approximate *P*_0_(*B*_*m-k*_; *B*_*k*_, *ζ*) ≈ *P*_0_(*B*_*m-k*_, *ζ*) and similarly, *P*_0_(*B*_*m-k*_; *B*_*k*_, *ζ*)*P*_0_(*B*_*k*−1_, *ζ*) ≈ *P*_0_(*B*_*m*−1_, *ζ*) to obtain:

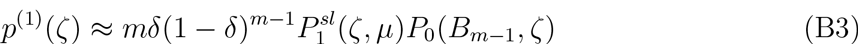

Here, we have ignored any correlations between successive B-Tfh cell interactions, and thus assumed that the order in which B cell interactions with self and non-self Tfh cells is unimportant. By making this approximation, we can generalize this to the case of *k* interactions between B cells and self reactive Tfh cells to obtain:

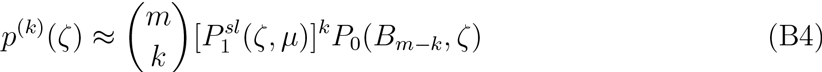

Eq. (B3) can be used to define a probability of m encounters between B cells and Tfh cells as:

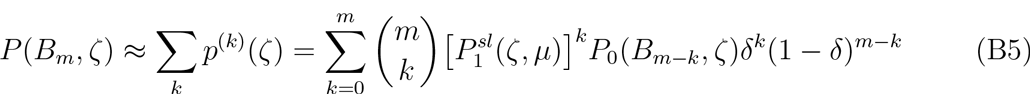

Since we have assumed that there is only one type of self recognizing Tfh cell present, the probability is a product of identical probabilities of single encounters with self Tfh cells, i.e 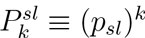. Thus, using the multiplicative property that *P*_0_(*B*_*m*_, *s*) ≈ *P*_0_(*B*_*m-k*_, *s*)*P*_0_(*B*_*k*_, *s*) we can divide Eq. (B5) by *P*_0_(*B*_*m*_, *ζ*) to obtain:

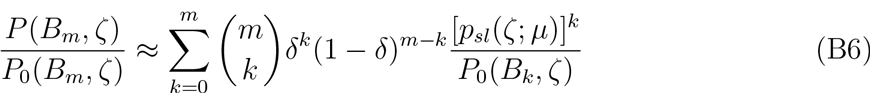

Define bounds on the probabilities of individual B-Tfh encountersnas:

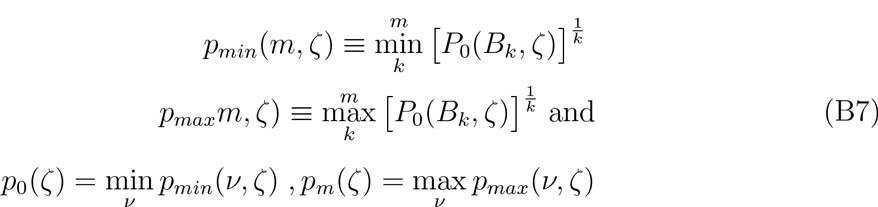

Physically, *p*_*min*_(*m*, *ζ*) represents a lower bound on the smallest effective probability of an individual B-Tfh cell encounter, and *p*_*max*_is the corresponding upper bound. Then, it is easily shown by summing the series, Eq. (B6) that:

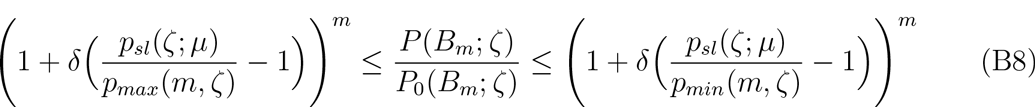

Furthermore, from the definitions Eq. (B6), we can write additional bounds on Eq. (B8) as

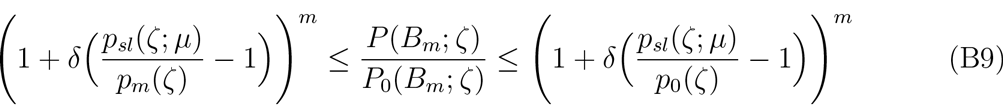

Eq. (B9) shows that when the probability of binding to a self Tfh cell is higher than the smallest effective probability, *p*_0_(*ζ*), of binding to a non-self Tfh cell, the presence of self Tfh cells creates a more favorable environment for affinity maturation of cross-reactive B cells. This is because when *p*_*sl*_(*ζ*,μ) > *p*_0_(ζ), the right hand side of Eq. (B9) becomes larger than 1, causing the probability of costimulation by m self reactive and antigen specific Tfh cell encounters to become greater than the probability of similar costimulation with only antigen specific Tfh cells. When antigen affinity is high towards both self recognizing and antigen specific Tfh cells, such that {*n*(*ζ*), *n*(*s*)} ≫ 1, the energy function in Eq.(18) can be approximated. To do so, consider that the energy function has the form:

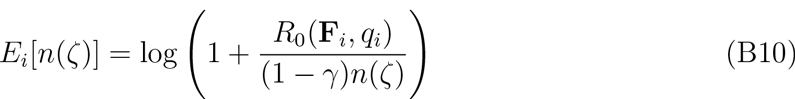

such that the ratio of survival probabilities between the cross-reactive B cell and an antigen specific B cell of high affinity becomes:

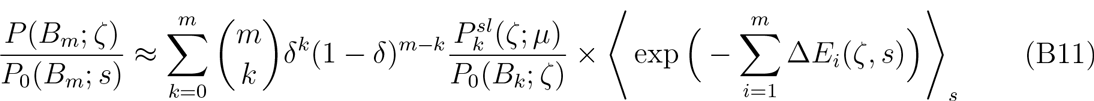

Combining Eq. (B11) with the inequalities, Eq. (B9), we obtain

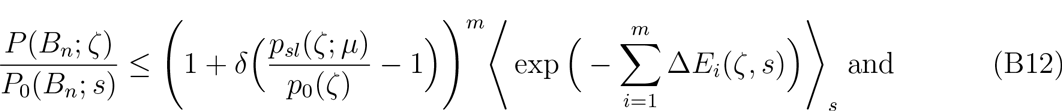

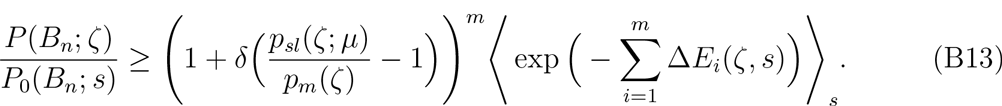

If we write 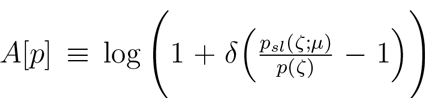 and 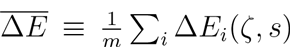 then the above inequalities have the form:

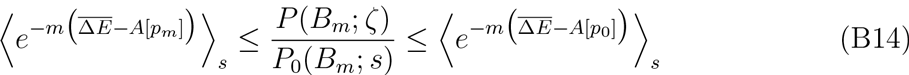

Thus, when the probabilities of self reactive encounters, *p*_*sl*_ are such that A[*p*_*m*_] > 0 and the average difference in ‘energies” 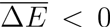 over the more probable sequences of B-Tfh cell encounters, we have a breakdown of tolerance, since the exponent on the left side of the inequality, Eq. (B14) is positive, and thus the probability of choosing autoreactive B cells over antigen specific B cells is exponentially higher. These conditions obtain when *p*_*sl*_(*ζ*, *μ*) > *p*_*m*_(*ζ*), i.e the probability of successful encounters with self reactive Tfh cells is greater than the maximal probability of encounters with antigen specific Tfh cells, in concert with the condition (1 − γ)*n*(*ζ*) ≳ *n*(*s*), i.e auto-reactive B cells also present comparable amount of antigen as the antigen-specific B cells. In brief this means that tolerance is broken when i) there are enough self reactive Tfh cells, and sufficient amounts of self antigens presented on autoreactive B cells so that such B-Tfh cell encounters are more favorable than corresponding encounters with antigen specific Tfh cells, and ii) the autoreactive B cells are also equally good at presenting non-self antigens, as purely antigen specific B cells.

## APPENDIX C: Probability distribution of number of clones

Consider a number *N* of *B* cells. Let the probability that a single B cell undergoes asymmetric cell division be *p* and probability that it does not undergo any division as *q* and the probability that it leaves the dark zone as *r*. Then, we can treat the clonal expansion of such B cells in the dark zone as a branching process. Since we are interested in the number of offspring only, we ignore the fact that each cell division results in a mutated BCR in one of the daughter cells. Such a branching process has a generating function;

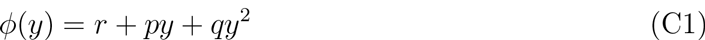

Correspondingly, let the number of clones after one generation for a single B cell be *Z*_0_, then the average number of progeny from a single B cell after a single generation is defined as 𝔼[*Z*_0_] = σ. It is easy to see that σ = *p* + *2q*. Let *Z*_*t*_ be a random variable describing the number of offspring after *t* generations from a single B cell. Then, we have the expected number of B cells after *t* generations is 𝔼[*Z*_*t*_] = σ^*t*^. The total number of clones, on average from a single B cell, in the dark zone after *T* generations is:

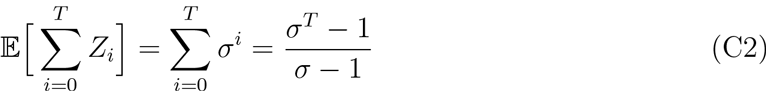

Consequently, if there are *N*(*B*_ν_, *s*) B cells present initially in the dark zone, we have the average number of B cell clones in the dark zone to be

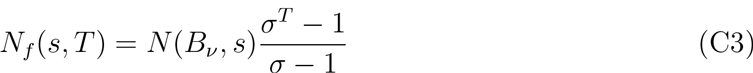

If the average time for a single generation of the B cell is τ, the growth rate can be defined as *γτ* ≡ log σ. Thus, we can write:

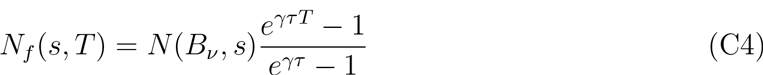

From experiments, we know that the duration of the cell cycle is related to the amount of costimulation of the B cell by Tfh cells, specifically for B cells with higher levels of costimulation, the duration of the cell cycle is shorter, or alternately, the growth rate of the B cells is longer. In addition, the total number of generations that the cells remain dividing for, increase with the levels of costimulation, Thus, if we define, *E*_*ν*_ as in the main text, we can write: *γ* ≡ *γ*(*E*_*ν*_, *s*) and *t*(*E*_*ν*_) ≡ *τ*(*s*)*T*(*s*). With these definitions we have

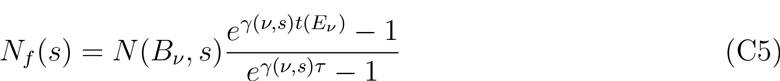

Substituting *N*(*B*_*ν*_, *s*) ≤ *N*(*B*_0_, *s*) exp(−*E*_*ν*_), we get

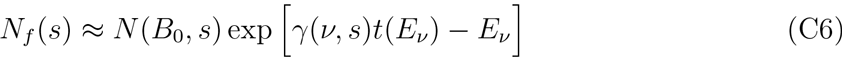

Here, we have used the property that the denominator, Eq. (C5) is of the order of unity, and also for long enough generations, the numerator is dominated by the exponential term.

